# Transvection and pairing of a *Drosophila* Hox long noncoding RNA in the regulation of *Sex combs reduced*

**DOI:** 10.1101/045617

**Authors:** Tom Pettini, Matthew Ronshaugen

## Abstract

Long noncoding RNAs have emerged as abundant and important regulators of gene expression in diverse animals. In *D. melanogaster* several lncRNAs involved in regulating Hox gene expression in the Bithorax Complex have been reported. However, no functional Hox long noncoding RNAs have been described in the Antennapedia Complex. Here we have characterized a long noncoding RNA *lincX* from the Antennapedia Complex, that is transcribed from previously identified *cis*-regulatory sequences of the Hox gene *Sex combs reduced (Scr)*. We use both the GAL4-UAS system and mutants to ectopically overexpress the *lincX* RNA from exogenous and endogenous loci respectively, in order to dissect the potential regulatory functions of *lincX* RNA versus *lincX* transcription. Our findings suggest that transcription through the *lincX* locus, but not the *lincX* RNA itself, may facilitate initiation of *Scr* in *cis* in the early embryo. Transvection phenomena, where regulatory sequences on one chromosome can affect expression on the homolog, have previously been reported in genetic studies of *Scr*. By analysing *lincX* and *Scr* nascent transcriptional sites in embryos heterozygous for a gain of function mutation, we directly visualize this transvection, and observe that the ectopic *lincX* transcriptional state appears to be relayed in *trans* to the homologous wild-type chromosome. This *trans*-activation of *lincX* correlates with both ectopic activation of *Scr* in *cis*, and increased chromosomal proximity. Our results are consistent with a model whereby early long noncoding RNA transcription through cis-regulatory sequences can be communicated between chromosomes, and facilitates long-range initiation of Hox gene expression in *cis*.

## INTRODUCTION

During animal development segmental identity is imparted through the function of highly conserved Hox transcription factors expressed in precise spatial and temporal patterns (McGinnis and Krumlauf 1992). In *Drosophila melanogaster*, Hox transcription occurs rapidly and is initially directed largely through the combined action of maternally deposited and early expressed transcription factors on *cis*-regulatory DNA elements (Harding and Levine 1988; Irish *et al*. 1989; Jack and McGinnis 1990). Interestingly, non-coding transcription is pervasive in the *cis*-regulatory dense intergenic regions of metazoan Hox complexes (Bae *et al*. 2002; Calhoun and Levine 2003; Rinn *et al*. 2007). Whereas a subset of this transcription may be spurious and arise from bi-valent or adventitious promoters, some of the transcripts have been shown to be conserved, and some have been demonstrated to be bona fide long noncoding RNAs (lncRNAs) with important functions in the regulation of Hox gene expression (Petruk *et al*. 2006; Rinn *et al*. 2007; Gummalla *et al*. 2012). Furthermore, the process of transcription alone can modulate the functions of some *cis*-regulatory elements (Rank *et al*. 2002; Ørom and Shiekhattar 2011; Lam *et al*. 2014). Thus it is now clear that many previously identified mutations resulting in homeotic transformations may disrupt the expression or potential functions of intergenic regulatory lncRNAs (Lipshitz *et al*. 1987; Sanchez-Herrero and Akam 1989; Pattatucci *et al*. 1991; Castelli-Gair *et al*. 1992). Understanding intergenic transcription and lncRNA function may be a key component in understanding the segment-specific activities of Hox regulatory DNA elements, and in turn the precise spatial and temporal control of Hox gene expression in development.

In the early 1950s Ed Lewis first began to dissect Hox regulatory interactions in the Bithorax Complex (BX-C) where he observed that chromosomal rearrangements affected the severity of homeotic phenotypes associated with a mutation on the other chromosome (Lewis 1954). He termed this communication of regulatory information between chromosomes the ‘transvection effect’ and it has since been genetically observed at many *Drosophila* Hox gene loci (Duncan 2002). Interestingly, the regulatory regions associated with Hox transvection in the BX-C are themselves transcribed. Previous genetic studies of the Hox gene *Sex combs reduced (Scr)* in the Antennapedia complex (ANT-C) revealed a distal regulatory region that contains multiple redundant positive and negative regulatory DNA elements (Gindhart *et al*. 1995). This region is also associated with numerous mutations resulting in a dominant gain of function (GOF) *Scr* homeotic transformation, whereby sex combs (normally specific to the male T1 leg) are ectopically formed on legs of the T2 and T3 segments (Pattatucci *et al*. 1991). Strikingly, five different GOF mutations associated with translocations or inversions with breakpoints mapping to the *fushi tarazu–Antennapedia (ftz-Antp)* interval result in mis-regulation of *Scr* in *trans*, on the wildtype (wt) chromosome (Southworth and Kennison 2002). Interestingly, these GOF mutations break within a region in the *ftz-Antp* interval where intergenic transcription has been previously observed (Calhoun and Levine 2003). One additional GOF mutation maps distal to this region in the first intron of *Antp*. Though transcripts from the *ftz-Antp* interval have been structurally determined by whole genome sequencing (Graveley *et al*. 2011) they have not been functionally characterized nor have the effects of these GOF mutations on transcription within the region been examined. We hypothesized that mutations in this region may affect intergenic transcription that is normally involved in regulating expression of the *Scr* gene in *cis* or in *trans*.

Here we have analysed the expression and function of a novel lncRNA we have named *lincX*, that overlaps previously characterized distal cis-regulatory elements of *Scr* in the *ftz-Antp* interval (Figure 1A). Transgenic and genetic overexpression approaches, as well as nascent transcript imaging were used to assess *lincX* functions both in *cis* and *trans*. Though we found no function for transgenically produced *lincX* RNA, *lincX* transcription was observed to be associated with activation of *Scr* in *cis*. Using nascent transcript FISH (ntFISH) for chromosome specific imaging in mutants, we found extensive ectopic *lincX* transcription in *trans*, a direct visualization of transvection. Transcription of ectopic *Scr* in *trans* was also observed, but almost always on chromosomes where *lincX* was also expressed. Further, we found that the transvection at *lincX* was associated with chromosomal pairing at or near the *lincX* locus.

**Figure 1.**
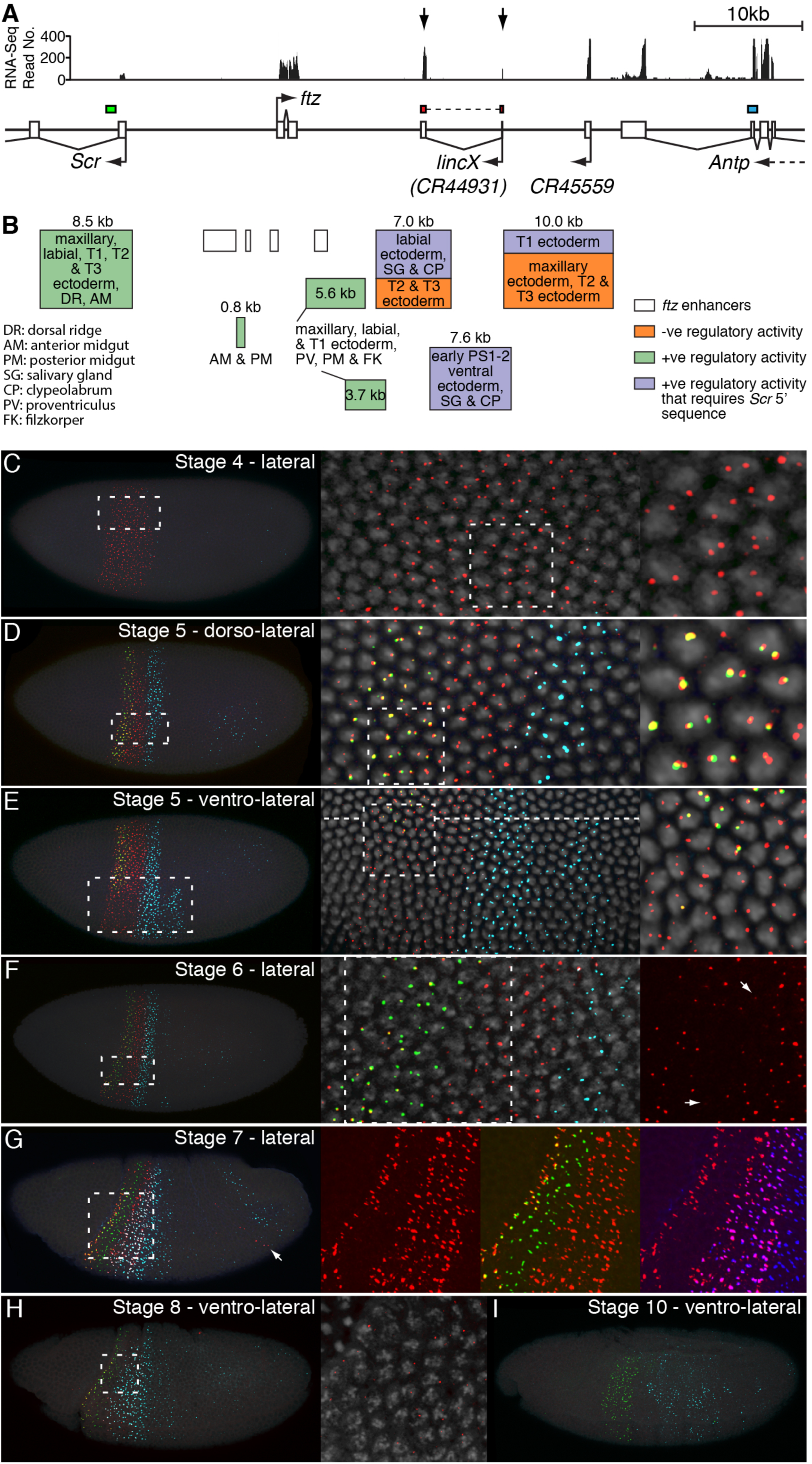
Transcription of lncRNA *lincX* is early and transient, preceding and overlapping *Scr* expression. A) The horizontal line represents genomic DNA in the *Scr-Antp* interval of the *D.melanogaster* ANT-C complex. Open boxes indicate exons, connecting lines introns. Colored bars above show positions of RNA probes used in ntFISH (see also Figure S2 probes 1, 4, 6). Top: RNA-seq data from 4-6 hour embryos (www.modencode.org). The 2 vertical arrows indicate 2 peaks of exonic transcription of lncRNA *lincX*. B) Previously identified regulatory elements in the *Scr-Antp* interval. The regulatory activity of orange, green and purple fragments was determined by Gindhart *et al*. 1995; the embryonic regions where expression is enhanced (green/purple) or repressed (orange) are indicated. C-I) ntFISH confocal images of *w*^*1118*^ *D.melanogaster* embryos; RNA probe positions are indicated in A); nuclei are counterstained with DAPI (grey), dashed boxes indicate the region magnified in right panels.

Transvection in the Hox complex is common, associated with lncRNAs, and may be a consequence of a gene regulatory mechanism normally meant to act over long distances in *cis* to stabilize interactions within a chromosome, but sometimes stabilizing interactions in *trans* between chromosomes. As regulation by lncRNAs is a common but poorly understood feature of Hox complexes in diverse animals, determining lncRNA function in *D.melanogaster* may provide important insight into the mechanisms of lncRNA function across phyla.

## MATERIALS AND METHODS

### *lincX* transcript structure and coding potential

Genomic sequence and automated annotations of the ANT-C were obtained from NCBI and transcriptome data from modENCODE. These were used to annotate splice junctions and approximate transcript ends for *lincX*. Multiple methods were used to assess *lincX* coding potential. See Extended Experimental Procedures for further details.

### Nascent transcript fluorescent in-situ hybridization (ntFISH)

RNA probe synthesis, embryo fixation, and ntFISH were performed according to (Kosman *et al*. 2004); see Extended Experimental Procedures for our specific adaptations to the protocols. Primer sequences for probe synthesis are provided in Table S1.

### Construction of *lincX* overexpression transgenics

Exonic *lincX* sequence was cloned in both orientations into expression plasmid pUAST, and constructs used in p-element-mediated transgenesis of the *w*^*1118*^ strain, performed by BestGene Inc. (Chino Hills, CA). Multiple independent transgenic lines were recovered for each construct (see Extended Experimental Procedures). Primer sequences for *lincX* cloning are provided in Table S1.

### Fly genetics

Full genotypes and genotype abbreviations of all fly lines used in this study are listed in Table S2, and details of all genetic crosses performed are provided in Table S3. Line maintenance and genetic crosses were performed at 25°C.

### *lincX* overexpression

Two independent transgenic lines for pUAST-lincX and pUAST-lincX-inverted constructs were crossed to Act5C-GAL4 and Dll-GAL4 driver lines. pUAST/GAL4 males were collected, and the number of sex comb teeth on each leg of males counted, as well as control genotypes +/GAL4 and *w*^*1118*^. ntFISH was used to visualise embryonic overexpression of transgenes (see Extended Experimental Procedures and Table S3).

### Mapping *Scr*^*W*^ inversion breakpoints

Fragments from around each *Sc*^*W*^ breakpoint were PCR-amplified from genomic DNA extracted from *Scr^4^Scr^W^*/TM3 adult flies, and sequenced (see Table S1 and Extended Experimental Procedures for primer sequences and combinations).

### Imaging and image analysis

Embryos were imaged using an Olympus FluoView FV1000 confocal laser scanning microscope and nascent transcriptional dots (nt-dots) were quantified manually. Standard light microscopy was used for imaging sex combs. For detailed imaging and quantification methods see Extended Experimental Procedures.

## RESULTS

### lncRNA *lincX* overlaps *cis*-regulatory sequences of the Hox gene *Scr*

The interval between the Hox genes *Scr* and *Antp* in the *Drosophila* Antennapedia Complex (ANT-C) contains the pair-rule gene *fushi tarazu (ftz)*, and two computationally annotated embryonically expressed long non-coding RNA transcripts: Dmel\CR44931 and Dmel\CR45559 (Figure 1A). The CR44931 transcript partially overlaps a transcript previously identified by Kuroiwa *et al*. 1985 originally named X; we have therefore renamed this transcript *long intergenic non-coding X (lincX)*. Using the genomic mapping of poly(A)+ RNA-seq reads available at http://modencode.org, we confirmed the annotated splice junctions and manually determined presumptive 5’ and 3’ ends for the *lincX* transcript (Extended Experimental Procedures). *lincX* has a 5’ exon of ~44bp and 3’ exon of ~351bp, separated by a 7096bp intron. Our annotation of *lincX* splice junctions exactly match those for transcript Dmel\CR44931, but our annotation of transcript ends is more conservative. We used a variety of methods to analyse the coding potential of the *lincX* RNA and found no evidence for coding potential (Extended Experimental Procedures).

Previous genetic and molecular studies identified multiple positive, negative, and redundant *cis*-regulatory elements responsible for controlling *Scr* expression (Gindhart *et al*. 1995) (Figure 1B). Several fragments were able to drive expression of a simple reporter gene in a subset of the normal embryonic *Scr* pattern (Figure 1B, green). Interestingly, three fragments from the *ftz-Antp* interval only drove reporter expression in combination with 2.3kb of sequence from immediately 5’ of the endogenous *Scr* promoter (Figure 1B, purple). These fragments were able to increase and refine aspects of normal *Scr* transcription, with two fragments, one of which overlaps *lincX*, also acting to repress expression outside of the wt *Scr* domain (Figure 1B, orange).

### *lincX* transcription precedes and fully overlaps expression of *Scr*

Multiplex nascent transcript FISH (mntFISH) was used to simultaneously determine the expression of *lincX, Scr* and *Antp* in *w*^*1118*^ *D.melanogaster* embryos (see Figure 1A, S2 and Extended Experimental Procedures for probe details). *lincX* expression initiates early and is first detected at stage 4, prior to the onset of *Scr* or *Antp* transcription at stage 5, in a stripe of even intensity both dorsally and ventrally that extends from the anterior boundary of parasegment 2 to midway through parasegment 4 (Figure 1C). At stage 5, transcription of *Scr* initiates in cells residing entirely within the broader *lincX* expression domain (Figure 1D) with *Scr* transcription initially weak or absent ventrally in the presumptive embryonic mesoderm (Figure 1E). Exonic *lincX* probes primarily detect RNAs accumulated at the site of *lincX* transcription (nt-dots) and very little/no signal is detectable elsewhere in the nucleus or cytoplasm, suggesting that *lincX* transcripts are either highly unstable or not effectively released from their site of synthesis. The overlapping *lincX* and *Scr* nt-dot signals (Figure 1D, yellow) may reflect the close linkage of *Scr* and *lincX* on the chromosome or possibly pairing. Interestingly, at stage 6, the size and intensity of *lincX* nt-dots weakens specifically in cells expressing *Scr* (Figure 1F, arrows) and by stage 7, *lincX* transcription is almost completely absent in *Scr*-expressing cells (Figure 1G). At stages 6 and 7, there is a weak transitory posterior domain of *lincX* expression in a band only ~1 cell wide (Figure 1G, arrow). All *lincX* expression becomes greatly reduced in stage 8 (Figure 1H) and is undetectable by stage 10 (Figure 1I). Our in-situ observations of temporal expression are supported by transcriptome data, which also indicates that *lincX* is unlikely to be maternally loaded and that it is not detected throughout the rest of development or in adults (Graveley *et al*. 2011) (Figure S1).

### *lincX* RNA transcribed from a transgenic locus does not affect *Scr* expression

To examine possible diffusible functions of *lincX* RNAs, we overexpressed mature *lincX* RNA in both sense and antisense orientation (pUAST-lincX, and pUAST-lincX-inverted) from ectopic locations using the GAL4:UAS system (Figure 2A), and assayed *Scr* expression and sex comb tooth number as a readout of *Scr* function (Extended Experimental Procedures). Transgenic UAS-lincX males were crossed to ubiquitous or tissue specific GAL4 driver virgin females (Act5C-GAL4 and Dll-GAL4 respectively). For each cross, pUAST/GAL4 males were recovered, and the number of sex comb teeth on all six legs counted and compared to control genotypes: *w*^*1118*^, +/Act5C-GAL4 and +/Dll-GAL4. No ectopic sex comb teeth were observed on T2 or T3 legs nor were any other phenotypes observed in male or female *lincX* overexpressing animals. Furthermore there was no significant change in T1 sex comb teeth number compared to the relevant +/GAL4 control (Figure 2B&C p>0.2, Mann-Whitney U).

**Figure 2.**
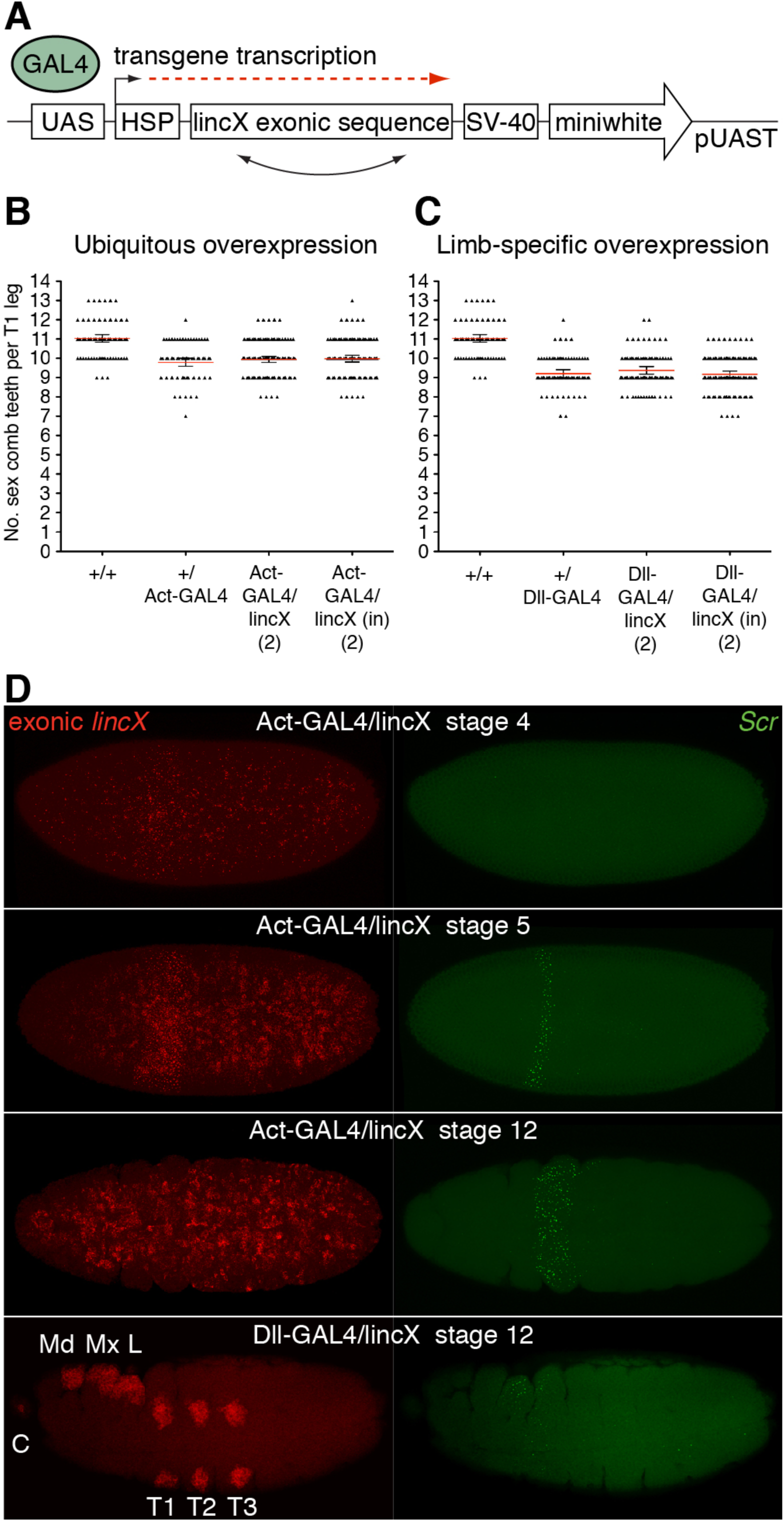
Mature *lincX* RNA does not function as a long range diffusible regulator of *Scr*. A) *lincX* exonic sequence was cloned in both orientations into pUAST inducible expression vector. UAS, GAL4 inducible upstream activating sequence; HSP, heat shock promoter; SV-40, termination sequence; miniwhite, eye color reporter. Transgenic lines were crossed to Act5C-GAL4 and Dll-GAL4 lines driving transcription through inserted *lincX* sequence. B-C) Quantification of T1 leg sex comb teeth (SCT) in males with Act5C-GAL4 (ubiquitous) and Dll-GAL4 (limb-specific) overexpression of pUAST-lincX and pUAST-lincX-inverted transgenes. The inverted (in) construct produces RNA antisense to endogenous *lincX*. Numbers in brackets indicate the number of independent transgenic lines, +, wt chromosome. Small black triangles indicate SCT number for each T1 leg; red line, mean; error bars are the 95% confidence interval. D) ntFISH confocal images of embryos with ectopic overexpression of a pUAST-lincX transgene induced by Act5C-GAL4 and Dll-GAL4 driver chromosomes. Anterior is left. Intronic *Scr* and exonic *lincX* probes are as in Figure 1A (also see Figure S2 probes 1&4). C, clypeolabrum; Md, mandibular; Mx, maxillary; L, labial; T1-T3, thoracic segments.

To determine if overexpression of *lincX* from an ectopic locus might affect *Scr* transcription in the absence of an observable phenotype, ntFISH was used to detect *Scr* and *lincX* accumulation in stage 4 to 14 embryos that expressed ectopic *lincX* (Figure 2D). Act5C-GAL4 driven overexpression of *lincX* RNA was first detected at stage 4, stochastically throughout the embryo. This coincides temporally with the onset of endogenous *lincX* transcription visible as a stripe of well-ordered nt-dots (Figure 2D). By stage 5, transgene expression was detectable in nearly all cells and throughout all later embryonic stages, though with an uneven level of expression among cells. Dll-GAL4 driven overexpression was first detected at stage 12 in the clypeolabrum, mandibular, maxillary, and labial segments, and in the limb primordia of T1-T3 segments. By this stage endogenous *lincX* is no longer expressed. We note that while cytoplasmic accumulation of the endogenous mature *lincX* RNA is minimal, the transgenic *lincX* RNA accumulates in both the cytoplasm and nucleus at appreciable levels. We propose that this may be a consequence of the stabilizing effect of the SV40 polyadenylation sequence. In stage 4 to 14 embryos we failed to observe any ectopic expression of *Scr*. This is consistent with our finding that overexpression did not have any phenotypic consequences, and suggests that the *lincX* RNA itself has no diffusible *trans* function on *Scr* transcription.

### A distant transposable element insertion in *Antp* causes ectopic *lincX* transcription from the endogenous locus

Overexpression of *lincX* from transgenic loci specifically tested for a diffusible function of the mature *lincX* RNA in regulation of *Scr*. However, to specifically examine the consequences of *lincX* transcription on *Scr* expression requires manipulation of *lincX* transcription from a structurally normal endogenous locus. Mutations mapping to the *ftz-Antp* interval are frequently associated with a homeotic *Scr* GOF phenotype where ectopic sex comb teeth form on T2 and T3 legs (Pattatucci *et al*. 1991; Southworth and Kennison 2002). However, one *Scr* GOF allele, *Antp*^*Scx*^, is a spontaneous ~3kb insertion of repetitive DNA (likely derived from a transposable element) that maps ~1.5kb downstream of the *Antp* P1 promoter (Hannah and Stromnaes 1955; Scott *et al*. 1983). This is more than 100kb from known *Scr* cis-regulatory regions, and more than 150kb from the *Scr* promoter (Figure 3A). We hypothesised that the *Antp*^*Scx*^ GOF sex comb phenotype may be associated with changes in *lincX* expression. To test this we performed ntFISH on *Antp*^*Scx*^/+ embryos, using probes shown in Figure 3A, and found that *lincX* is expressed in both its normal domain and a posterior ectopic domain precisely overlapping the *Antp* P2 pattern (compare Figure 3B&C). Probes against *Antp* P1 and *Antp* P2 promoters, and a common *Antp* 3’ probe were used across a developmental time course of *w*^*1118*^ embryos to identify the specific pattern (Figure S3).

**Figure 3.**
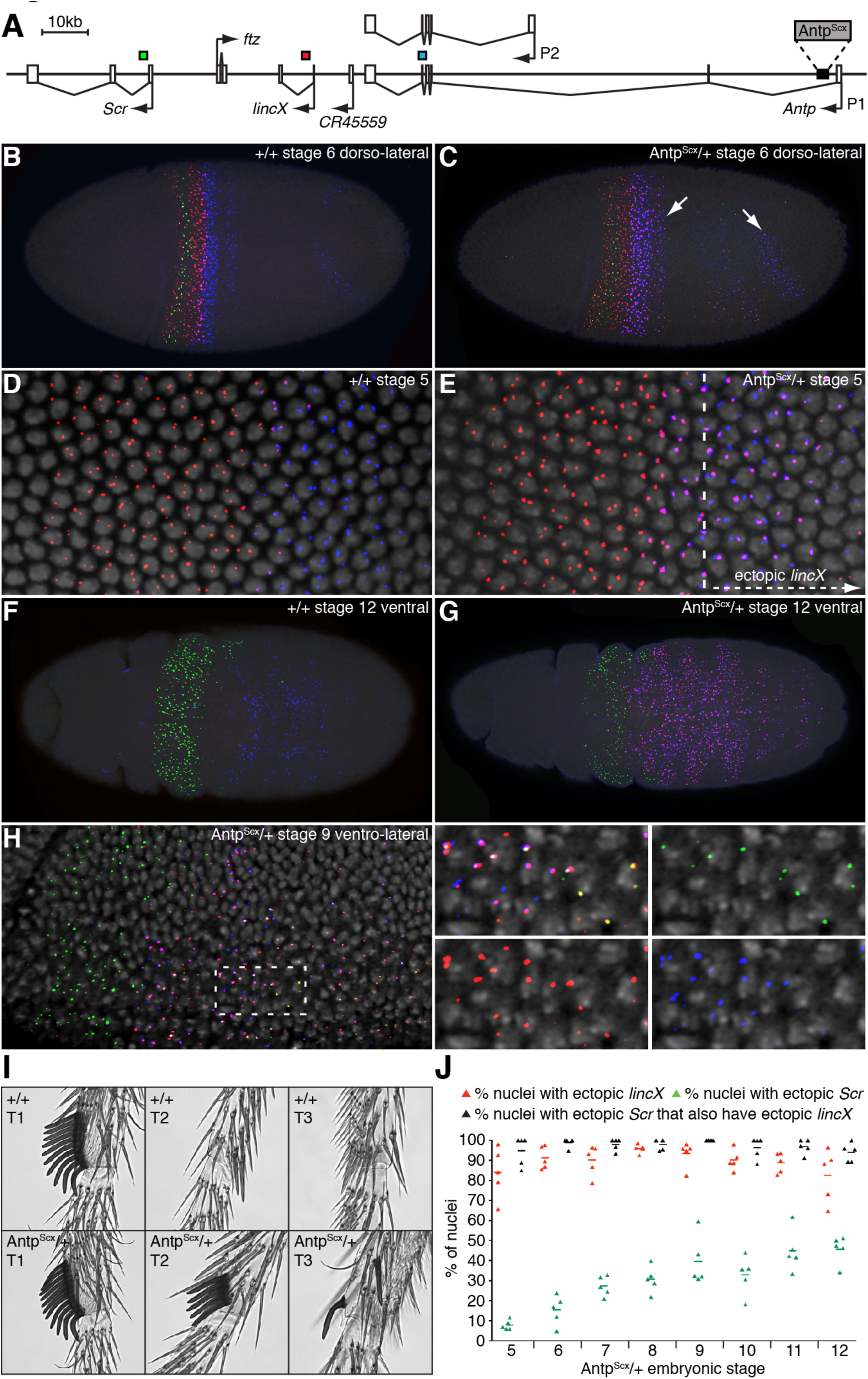
The *Antp*^*Scx*^ mutation causes ectopic *lincX* and *Scr* expression and results in a GOF *Scr* phenotype. A) Schematic showing the ~1.5kb *Antp*^*Scx*^ retro-transposon insertion. *Scr, lincX* and *Antp* probe positions used for ntFISH experiments (B-H) are shown as colored bars (see also Figure S2 probes 1, 3, 6). B-H) ntFISH on *w*^*1118*^ and *Antp*^*Scx*^/+ heterozygous embryos, colors correspond to probes shown in A). Overlap between *lincX* and *Antp* P2 signals is visible as purple. C) Arrows highlight domains of ectopic *lincX* transcription. E) Vertical dashed line is a conservative estimate of the posterior boundary of wt *lincX* expression. H) Dashed box is magnified to the right. I) Representative images of the presumptive sex comb forming regions in male T1-T3 legs. The *Antp*^*Scx*^ mutation causes a GOF ectopic sex comb phenotype. J) Quantification of ectopic *lincX* and *Scr* expression over developmental time in *Antp*^*Scx*^/+ embryos. Plot shows % of sampled nuclei with ectopic *lincX* transcription (red), ectopic *Scr* (green) and the % of those nuclei with ectopic *Scr* that also had ectopic *lincX* (black). Each point represents 1 embryo, in almost all embryos >50 nuclei were sampled. See also Figure S3.

ntFISH allows us to resolve the transcriptional activities on the wt and mutant *Antp*^*Scx*^ chromosomes in individual nuclei. In wt stage 5 embryos *lincX* expression overlaps the *Antp* P2 domain by only ~2 cells (Figure 3D, pink overlap). In Figure 3E, a dotted line marks the ectopic expression boundary in an *Antp*^*Scx*^/+ stage 5 embryo. To the left of this line, in the wt domain, nuclei express *lincX* from both chromosomes. To the right, specifically in the *Antp* P2 domain, ectopic *lincX* is expressed predominantly from only one chromosome. We infer this to be the mutant *Antp*^*Scx*^ chromosome. We have performed ntFISH in *Antp*^*Scx*^/+ embryos against transcript CR45559, which lies between *Antp* and *lincX* (Figure 3A) and found no ectopic expression, arguing against transcriptional read-through from *Antp* (data not shown). Instead we propose that on the *Antp*^*Scx*^ chromosome sequences that normally direct *Antp* P2 expression now also drive *lincX* transcription. This does not appear to disrupt normal *Antp* P2 expression, which is still transcribed from both chromosomes (Figure 3E). After stage 9, both wt and *Antp*^*Scx*^/+ embryos no longer express *lincX* in the endogenous domain (Figure 1I, 3F). However in *Antp*^*Scx*^/+ mutants, ectopic *lincX* expression continues throughout later development in the *Antp* P2 domain (compare Figure 3F&G).

### Ectopic transcription of *lincX* and *Scr* are correlated

In the domain of ectopic *lincX* transcription we also frequently observed ectopic *Scr* transcription (Figure 3H). This is consistent with the ectopic T2 & T3 sex comb phenotype observed in mutant adults (Hannah and Stromnaes 1955) (Figure 3I). Interestingly ectopic *Scr* nt-dots appeared to increase in frequency through embryogenesis and to arise primarily on chromosomes transcribing ectopic *lincX* (Figure 3H). To quantify this, nuclei in the *Antp* P2 domain in stage 5-12 *Antp*^*Scx*^/+ embryos were scored for the number and arrangement of *Scr, lincX* and *Antp* nt-dots (Figure 3J and see Extended Experimental Procedures). At all time points ectopic *lincX* was transcribed in ~90% of cells, whereas ectopic *Scr* was initially transcribed in only ~5% but steadily increased to ~50% by stage 12. Virtually all cells with ectopic *Scr* also expressed ectopic *lincX* (Figure 3J, black points). If ectopic activation of *Scr* and *lincX* were independent of one another, then in cells ectopically expressing *Scr* we would expect to find ectopic *lincX* at the same frequency as the overall proportion of cells with ectopic *lincX*. In fact, we found that across eight time points and 40 embryos, in 39 of the 40 embryos the percentage of ectopic *Scr* expressing cells also expressing ectopic *lincX* was higher than the observed overall percentage of ectopic *lincX* expressing cells in that embryo. In other words, ectopic *Scr* was expressed more frequently in *lincX* expressing cells than expected. These results are largely correlative and may indicate only active chromatin marks or indiscriminate long-range enhancer promoter interactions at two closely linked genes. However, the results are also consistent with a mechanism whereby *lincX* transcription facilitates subsequent initiation of *Scr* transcription.

A different *Scr* GOF mutation, *Scr*^*W*^ is caused by a ~50kb inversion with breakpoints within the *lincX* intron, and just 3’ of *Antp* P2 (Figure S4A). We found that this inversion allows transcriptional read-through from *Antp* P1, through the inverted non-coding transcript CR45559, and into the non-inverted 3’ end of *lincX* (Figure S4C). Thus, like *Antp*^*Scx*^*, Scr*^*W*^ also causes late *lincX* transcription in an ectopic posterior domain, but via the process of transcriptional read-through rather than the apparent indiscriminate enhancer activity of *Antp*^*Scx*^. This is consistent with a mechanism whereby *lincX* transcription facilitates initiation of *Scr* transcription, as we also observed ectopic transcription of *Scr*, co-expressed with ectopic *lincX*, in *Scr*^*4*^*Scr*^*W*^/+ embryos (Figure S4D).

### Visualization of transvection at the *lincX* and *Scr* loci

Analysis of nascent transcriptional sites in *Antp*^*Scx*^/+ embryos revealed that ectopic *lincX* was also transcribed from the wild-type chromosome (Figure 4A&B). This is contrary to the expectation that in a heterozygote only expression from the mutant chromosome should be affected, and suggests communication of regulatory information between the homologs. Our direct molecular observation of this ‘transvection’ is consistent with previous genetic observation of transvection associated with *Antp*^*Scx*^ (Southworth and Kennison 2002). To quantify this transactivation effect, nuclei from the ectopic *lincX* expression domain in *Antp*^*Scx*^/+ embryos were sampled across stages 5-12 as described above (see also Extended Experimental Procedures). The proportion of nuclei showing *trans* expression of ectopic *lincX* increased from stages 5 to 8 (~15% to ~40%, Figure 4C). After stage 8 *trans-lincX* expression was observed consistently in ~35-40% of nuclei. Transactivation of *Scr* was less frequent and hardly observed before stage 8 (3%), but increased throughout embryogenesis reaching 12% by stage 12 (Figure 4C). We focused on nuclei showing *trans*-activation of *Scr* (2 *Scr* nt-dots) and asked what proportion also had *trans*-activated *lincX*. If their expression were independent the expected proportion would simply be equal to the percentage of *trans-lincX* nuclei, about 40%. Strikingly, this proportion was consistently around 70%, a marked enrichment in the association of *trans-Scr* with *trans-lincX* on otherwise wt chromosomes. Again, this correlation may reflect the linkage of *Scr* and *lincX* on the chromosome. However, the result is also consistent with *lincX* transcription itself facilitating activation of *Scr* primarily in *cis*, as ectopic *Scr* expression in *trans* is rare on chromosomes that do not express *lincX*. Furthermore this agrees with the transgenic *lincX* overexpression results above that suggested *lincX* RNA is unlikely to have a diffusible *trans* function.

**Figure 4.**
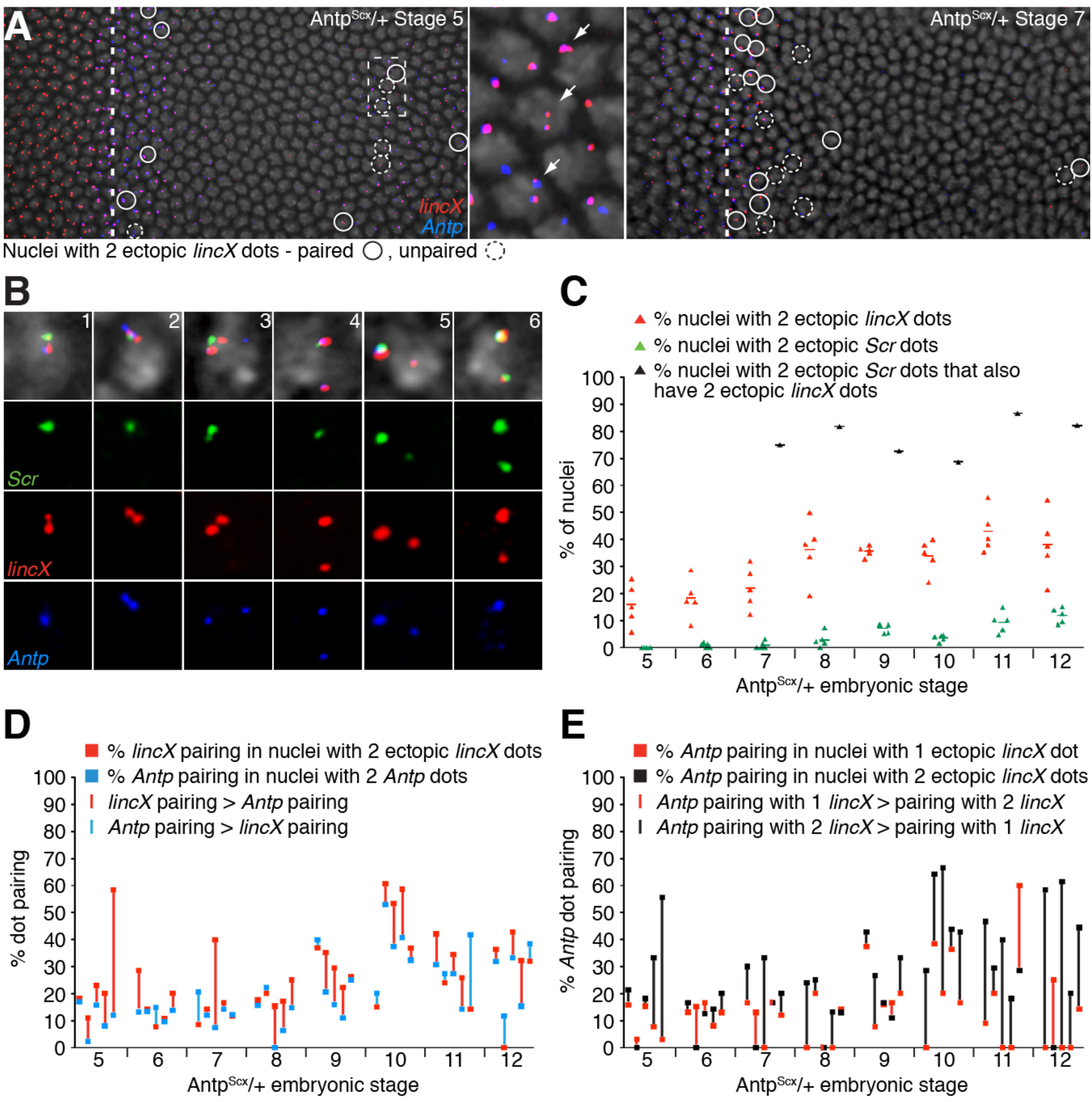
The *Antp*^*Scx*^ mutation induces ectopic *lincX* and *Scr* transcription in *trans* on a wt chromosome. A) ntFISH on *Antp*^*Scx*^/+ heterozygous embryos at stages 5&7, colors correspond to probes shown in Figure 3A, nuclei are counterstained with DAPI (grey). The vertical dashed line is a conservative estimate of the posterior boundary of wt *lincX* expression; dashed box is magnified in centre. Nuclei with 2 ectopic *lincX* nt-dots (indicating ectopic activation on the + chromosome in *trans)* are circled, and indicated by arrows in the magnified box. Dashed circles indicate unpaired *lincX* nt-dots and solid circles indicate paired *lincX* nt-dots. B) Nuclei from *Antp*^*Scx*^/+ heterozygous embryos, showing representative nt-dot arrangements of ectopic *Scr* and ectopic *trans*-activated *lincX*. C) Quantification of ectopic *lincX* and *Scr trans*-activation over developmental time in *Antp*^*Scx*^/+ heterozygous embryos. Plots show % of sampled nuclei with *trans-lincX* (red), *trans-Scr* (green) and % of *trans-Scr* nuclei that also had *trans-lincX* (black). Each red or green point represents 1 embryo, in almost all embryos >50 nuclei were sampled. Due to low incidence of *trans-Scr*, no black point is shown for stages 5 and 6. Black points do not have error bars as data was pooled across 5 embryos per stage. D) Frequency of pairing between ectopic *lincX* nt-dots (red points) and between *Antp* nt-dots (blue points) in *Antp*^*Scx*^/+ heterozygous embryos. Vertically aligned red and blue points are plotted for each embryo sampled with the connecting line color indicating whether frequency of pairing was higher for *lincX* (red) or *Antp* (blue). E) Frequency of pairing between *Antp* nt-dots in nuclei with 1 (red points) or 2 (black points) ectopic *lincX* nt-dots in *Antp*^*Scx*^/+ heterozygous embryos. Vertically aligned red and black points are plotted for each embryo sampled with the connecting line color indicating whether frequency of *Antp* nt-dot pairing was higher in nuclei with only 1 ectopic *lincX* (red) or with 2 ectopic *lincX* (black) nt-dots.

The *trans*-activation of *lincX* and *Scr* in *Antp*^*Scx*^/+ embryos may arise from indiscriminate long-range interactions with the *Antp* P2 enhancer in both *cis* and *trans* (Figure 5A). Alternatively, initial ectopic activation of *lincX* on the mutant chromosome in *cis* may subsequently lead to a relay of the ectopic transcriptional state in *trans* to the wild-type chromosome (Figure 5B). Observations from the *Scr*^*W*^ mutant lend support to the latter model. We found that in *Scr*^*4*^*Scr*^*W*^/+ embryos, *lincX* also frequently exhibited transvection, with ectopic transcription from both the mutant and wild-type chromosomes (Figure S4D). Less frequently *trans*-activation of *Scr* was also observed, consistent with the observed ectopic sex comb phenotype in *Scr*^*4*^*Scr*^*W*^/+ adults, as *Scr*^*4*^ is a protein null (Southworth and Kennison 2002). Since transcriptional read-through underlies the ectopic *lincX* transcription in the *Scr*^*W*^ mutant, it is unclear how this cis-specific mechanism alone could lead to ectopic transcription on the homolog. A mechanism whereby initial ectopic *lincX* transcription on one chromosome can itself activate *lincX* transcription from the copy in *trans* could help explain the observation.

**Figure 5.**
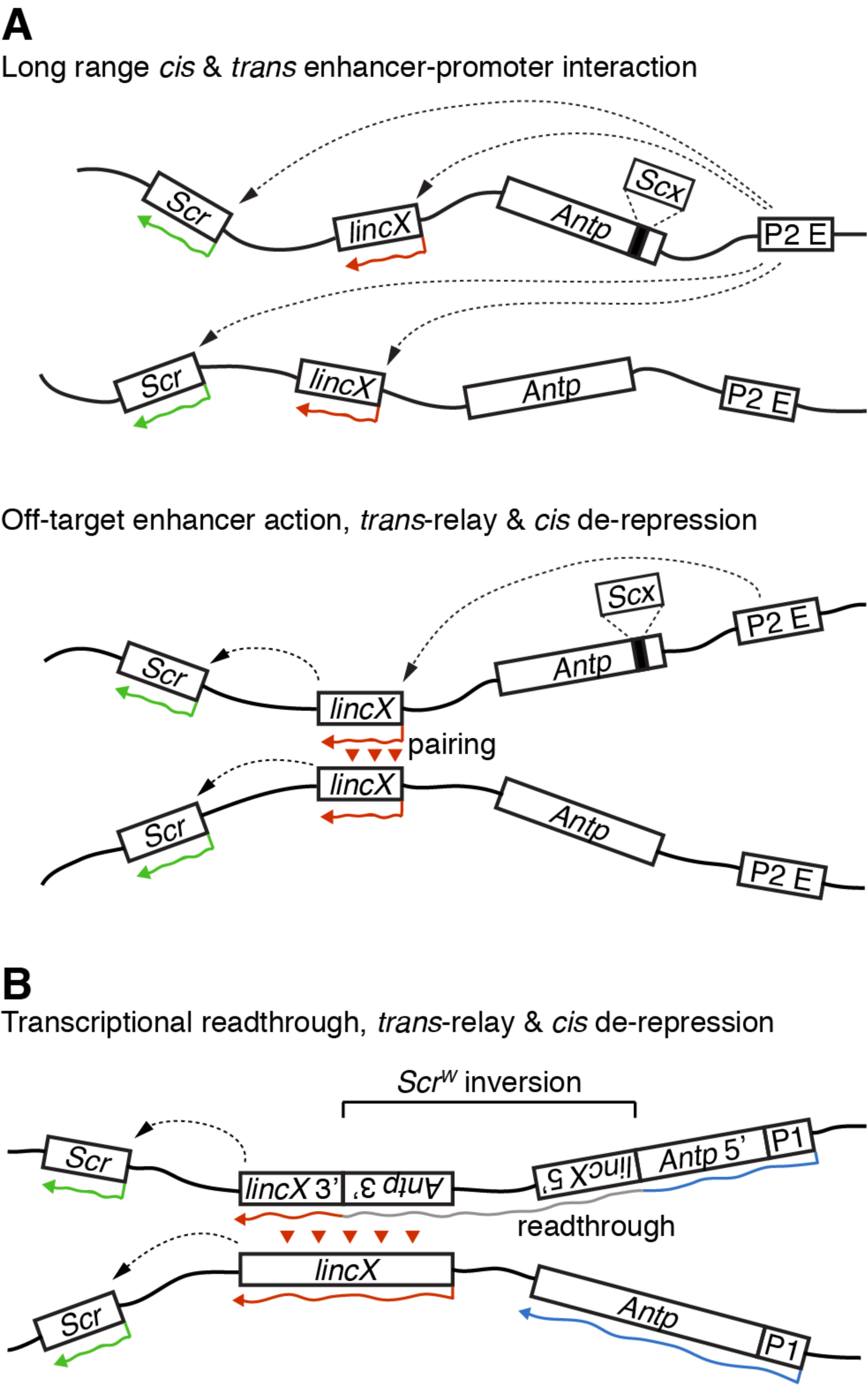
Models of ectopic *cis/trans* transcription of *lincX* and *Scr* in *Antp*^*Scx*^ and *Scr*^*W*^ mutants. A&B) Dashed arrow lines represent activation or de-repression, solid coloured arrows represent active transcription, red arrow heads indicate a relay of transcriptional state in *trans* between paired *lincX* loci. A) Two alternative models of ectopic *lincX* and *Scr* transcription in *cis/trans* in an *Antp*^*Scx*^/+ heterozygous embryo. P2 E is the *Antp* P2 enhancer. B) A model of ectopic *lincX* and *Scr* transcription in *cis/trans* in an *Scr*^*W*^/+ heterozygous embryo. The grey portion of the arrow represents transcription through inverted sequence, producing a transcript that is not normally made from a wild-type chromosome.

### The *lincX* locus is near a focal point for chromosome pairing

In *Antp*^*Scx*^/+ embryos, we observed that in nuclei showing *lincX* transvection, frequently the two *lincX* nt-dots were paired (defined as overlap of flourescence signal with separation <400 nm), suggesting physical proximity of the two chromosomes (Figure 4A & 4B panels 1-3). To determine if chromosome pairing is broadly distributed across the ANT-C or more localized near the *lincX* locus, we compared the pairing at the 5’ end of *lincX* with the 3’ end of *Antp* (separated by ~25kb). Nuclei with two ectopic *lincX* nt-dots (transvection) or two *Antp* 3’ nt-dots (wt) were sampled from the same region in stage 5-12 *Antp*^*Scx*^/+ embryos, and pairing was scored. For individual embryos we plotted the pairing percentage for *lincX* and *Antp* as two points connected by a colored vertical line indicating whether *lincX* pairing (red) or *Antp* pairing (blue) was more frequent (Figure 4D). Across all stages, in 30 out of 40 embryos sampled, pairing at the *lincX* locus occurred more frequently than at the *Antp* locus, χ^2^ (1, N = 40) = 10, p<0.002. This difference is dramatic given how close the two regions are on the chromosome and indicates that the *lincX* locus may be at or near a focal site for physical pairing between the homologs.

### Chromosome pairing and transvection of *lincX* are correlated

To investigate the association between chromosome pairing and transvection, we assayed *lincX* transcription and used a probe against the 3’ end of the *Antp* locus as a conservative proxy for nearby pairing. In stage 5-12 *Antp*^*Scx*^/+ embryos we scored pairing at *Antp* when either one (red point) or both (black point) *lincX* loci were ectopically transcribed (Figure 4E). For each embryo the color of vertical line connecting the points indicates if *Antp* pairing was higher in nuclei transcribing *lincX* from one (red) or both (black) chromosomes. Across all stages, in 30 out of 40 embryos sampled the incidence of pairing was higher in nuclei transcribing *lincX* from both chromosomes, showing a significant association between pairing and transcription, χ^2^ (1, N = 40) = 10, p<0.002. We can interpret this result in two different ways: it either indicates that *lincX* transcription from both chromosomes enhances physical pairing between the homologs at or near the *lincX* locus, or that physical proximity of the chromosomes enhances the *trans-activation* of *lincX*.

## DISCUSSION

It has been known for nearly 30 years that the *Drosophila* BX-C contains a number of lncRNAs interlaced within the complex regulatory regions that direct Hox expression (Lipshitz *et al*. 1987; Bae *et al*. 2002; Gummalla *et al*. 2012). Recently, human and mouse Hox complexes have also been found to contain numerous lncRNAs (Rinn *et al*. 2007; Delpretti *et al*. 2013). Some function in regulating Hox gene transcription in *cis*, some act as diffusible regulators in *trans*, and many work through interactions with chromatin modifiying proteins such as PRC2 (Rinn *et al*. 2007; Dinger *et al*. 2008; Tsai *et al*. 2010; Wang *et al*. 2011; Delpretti *et al*. 2013). We have characterized a lncRNA in the ANT-C transcribed through a region critical for the regulation of *Scr*. Though a variety of models have been proposed for lncRNA functions, dissecting lncRNA functions from those of the underlying regulatory DNA has been difficult. Here, we have used GOF transgenics, and taken advantage of a mutation that alters lncRNA expression without altering the sequence of the DNA at the endogenous locus, to discriminate between these functions. Furthermore, multiplex ntFISH allowed the simultaneous examination of lncRNA and Hox transcription on individual chromosomes, facilitating dissection of the *cis* and *trans lincX* activities.

In *Drosophila*, initial patterns of Hox expression are established rapidly in the first three hours of development following the maternal to zygotic transition (MZT). We find that *lincX* transcription initiates earlier than nearby Hox genes *Scr* and *Antp*. This is consistent with its identification as a direct target of Zelda, a transcription factor involved in activation of the earliest genes expressed at the MZT (Fu *et al*. 2014). Most other Hox lncRNAs such as *bxd* upstream of *Ubx* are also observed to be Zelda targets that are expressed very early in embryogenesis (Petruk *et al*. 2006; Fu *et al*. 2014). Early lncRNA expression may reflect the actions of pioneer transcription factors that open chromatin for access by other early transcription factors such as those encoded by the gap genes, establishing a prepattern of accessible and inaccessible chromatin and modulating the activities of regulatory elements.

Overexpression of the mature *lincX* RNA from ectopic locations had no observable effect on *Scr* transcription or any observable phenotypic consequence. However, two distinct chromosomal aberations result in ectopic transcription of the endogenous *lincX* locus that is correlated with ectopic activation of *Scr* in *cis*. Taken together, these results are consistent with a model where early embryonic transcription of *lincX*, but not the *lincX* RNA itself, facilitates initiation of *Scr* transcription in *cis*. We suggest that ectopic *lincX* transcription may attenuate the activity of the underlying T2-T3 repressive cis-regulatory DNA element within the *lincX* locus, thereby removing repression of *Scr* rather than directly activating it (Gindhart *et al*. 1995). Other Hox lncRNAs including *bxd* and iab-8 ncRNA have also been suggested to function in *cis* at the level of their transcription, but attenuate expression of their adjacent Hox genes via transcriptional interference (Petruk *et al*. 2006; Gummalla *et al*. 2012). Furthermore, recent examples have emerged where transcription modulates enhancer function (Lam *et al*. 2014). Thus the presence of transcription factors within a cell is not necessarily the ultimate arbiter of *cis*-regulatory element function as transcription through an element can be an additional mechanism for achieving different segment-specific activities.

Using ntFISH we were able to visualize transcription states on individual chromosomes in wild-type and mutant embryos. Surprisingly we found ectopic activation of *lincX* occurred not only in *cis* but also in *trans* on the wt chromosomes. In the *Antp*^*Scx*^/+ mutant, a transposable element insertion 120kb away from *lincX* in the first intron of *Antp* likely enables an *Antp* P2 enhancer to ectopically activate *lincX* transcription in the *Antp* P2 domain with almost perfect fidelity on the *Antp*^*Scx*^ chromosome. It is possible therefore that ectopic *lincX* expression, in *trans*, on a wildtype chromosome reflects an extreme form of this long-range enhancer-promoter interaction (Figure 5A, top). Alternatively, the transcriptional state of the *lincX* locus on the mutant chromosome may be communicated more directly if there is pairing at the locus that can subsequently induce activation of *lincX* in *trans* on the homolog (Figure 5A, bottom). In these models, the ectopic transcription of *Scr* may either result from direct long-distance enhancer-promoter interactions in *cis* and *trans* (Figure 5A, top), or alternatively from transcription of *lincX* in *cis* (Figure 5A, bottom). The latter is consistent with the observed strong association of *trans-Scr* with *trans-lincX* on otherwise wt chromosomes. Given the recent advances in precise genome editing via the CRISPR-Cas9 system, it may be possible to discriminate between the two alternate models proposed in Figure 5A. Specifically, deleting the *lincX* promoter on the *Antp*^*Scx*^ chromosome would allow us to test whether *lincX* transcription in *cis* is a prerequisite for subsequent ectopic activation of *lincX* (and *Scr)* in *trans* on the wild-type chromosome.

In contrast to *Antp*^*Scx*^, where a distant regulatory element directs the ectopic expression, the ectopic *lincX* transcription observed on the *Scr*^*4*^*Sc*^*W*^ mutant chromosome is caused by the cis-specific mechanism of transcriptional read-through. It is unclear if or how transcriptional read-through of the *lincX* region could contribute to *lincX* or *Scr* activation in *trans*. One possibility is that the *Sc*^*W*^ inversion disrupts the activity of a *trans*-acting repressive DNA element. However, this seems unlikely to be the cause, given that no ectopic sex comb teeth are formed in *Df(3R)Scx*^*4*^*/TM3* males, where *Df(3R)Scx*^*4*^ deletes a large region between the *lincX* intron and 5’ of *Antp* (Pattatucci *et al*. 1991). An alternative explanation is the same as one suggested for *Antp*^*Scx*^; that the transcriptional state of *lincX* on the mutant *Scr*^*W*^ chromosome induces ectopic transcription of *lincX* on the wild-type homolog through pairing (Figure 5B).

Interestingly, even though ectopic *Scr* transcription was predominantly observed in *cis* on the *Scr*^*W*^ mutant chromosome, genetic evidence suggests that only Scr protein produced in *trans* on the wild-type chromosome contributes to the adult ectopic sex comb phenotype. This follows from the observation that when the *Scr*^*4*^ LOF mutation is in *trans* to the *Sc*^*W*^ mutation the phenotype is almost completely supressed (Southworth and Kennison 2002). This discrepancy between early expression and adult phenotype suggests that the early *cis* expression does not reflect later *Scr* production and that the early initiation of *Scr* expression is distinct from its subsequent maintenence in the imaginal discs. Perhaps the continued expression of *Scr*, including in the imaginal discs, requires the action of a maintenance element whose function is disrupted by the inversion on the mutant chomosome, but remains intact and functional on the wild-type chromosome.

Recent studies have shown that transvection effects are a relatively common phenomenon throughout the *Drosophila* genome (Mellert and Truman 2012). Though transvection at the *Scr* locus was previously deduced from genetic studies (Pattatucci and Kaufman 1991; Southworth and Kennison 2002), here we have directly visualized the transcriptional state of both chromosomes at a transvecting locus, providing an important insight into the actual transcriptional profile on the mutant and transvecting wild-type chromosome through the early stages of a developmental process.

By analyzing nt-dot organization, we observed physical pairing of the *lincX* loci more frequently than pairing at a relatively nearby position (~25 kb away) in *Antp*, suggesting that *lincX* may be at or near to a focal point for *trans*-chromosome interactions. Furthermore, we revealed an association between *lincX* transcription in *trans*, and this chromosomal pairing. This finding is consistent with previous observations showing that transvection effects at the *Abd-B* locus involve pairing (Ronshaugen and Levine 2004). Southworth and Kennison 2002 showed that *Scr* regulatory sequences preferentially act in *cis*, but can act in *trans* when the *Scr* promoter in *cis* is absent. This *trans*-chromosome communication of regulatory information is common throughout the Hox complex of *Drosophila* not only at *Scr*, but also *Ubx, abdA*, and *AbdB* (Duncan 2002). Intriguingly, each of these Hox genes have associated adjacent regulatory intergenic lncRNAs; *bxd* upstream of *Ubx*, and a complex ncRNA rich region between *abd-A* and *Abd-B* containing *Mcp, Fab7* and iab-8 ncRNA (Rank *et al*. 2002; Petruk *et al*. 2006; Gummalla *et al*. 2012). Furthermore, the *yellow* locus which for many years has been studied as a classic example of a transvecting locus has recently been shown to be located immediately adjacent to a regulatory lncRNA called *yellow-achaete intergenic RNA (yar)* (Soshnev *et al*. 2011). This leads to the intriguing possibility that lncRNA functions may somehow be associated with transvection effects not only at these loci but more generally.

Given the role that both tandem and whole genome duplication has played in the genesis of the Hox complex coupled with the stunning conservation of syntenic and transcriptional Hox organization, it is an exciting possibility that lncRNA mediated regulation of Hox expression may be an ancestral feature of Hox regulation and that these units might have duplicated and diversified from an ancestral lncRNA mediating gene expression in *cis* and *trans*.

## ACKNOWLEDGEMENTS

This work was supported by The Manchester Fellowship to MR. Work on this project benefited from the Faculty's Fly Facility, established through funds from University and the Wellcome Trust (087742/Z/08/Z). We are grateful to the Papalopulu lab for generously sharing microscopy equipment, to Sanjai Patel who manages the Fly Facility, and we thank Fiona He and Janet Melling for technical assistance. We also thank Kimberly Mace and Sam Griffiths-Jones for their helpful suggestions.

